# Multi-Domain Characterization and Rapid Detection Technology for Cocaine and Methamphetamine Acute Abuse based on EEG

**DOI:** 10.1101/2022.07.21.500973

**Authors:** Jie Xia, Jintao Wu, Jiadong Pan, Fan Zhang, Hao Jin, Shurong Dong, Yueming Wang, Jikui Luo, Ke Si

**Author notes:** Jie Xia and Jintao Wu contributed equally to this work.

## Abstract

Multi-domain characterization and a new fast detection method for acute illegal psychostimulants abuse detection based on waking-state EEG of mice is proposed in this paper. To get corftical electroencephalogram (EEG), three groups of mice were injected with cocaine (Coca), methamphetamine (Meth), and saline (Sali) respectively following the experimental paradigm of drug abuse. The EEG features were extracted out by multi-domain views, including time, frequency, complexity, dynamics, and independent domains to obtain acute drug abuse effects on the brain. New strategy combing time domain with frequency domain is developed as multi-domain input and by means of dimension transformation approach preserving temporal information, the performance of drug abuse detection is greatly improved with deep learning models of ResNet50. Results show that comparing to support vector machine (SVM), k nearest neighbor (kNN), random forest (RF), and long short-term memory fully convolutional networks (LSTM-FCN), ResNet50 based on our proposed multi-domain features has best F1-score of 95.10%. This promising method provides a low-cost, fast, and widely assisted technology for psychostimulants abuse identification.

## 1 Introduction

Psychostimulants are the most commonly used psychoactive substances in the world (Gannon, Reichard, & Fantegrossi, 2014). They cause arousal and high mood, and increase alertness and arousal (Favrod-Coune & Broers, 2010). The abuse of illegal psychostimulants can lead to psychological dysfunction and behavioral disorders (Blume, 2006), which further seriously affect public health and social problems, such as human immunodeficiency virus (HIV) (Gannon et al., 2014) infection and the spread of COVID-19 (Schlosser & Harris, 2020). Cocaine (Coca) (Favrod-Coune & Broers, 2010) and methamphetamine (Meth) (Gock & Skrinska, 2005) are the most widely used illegal psychostimulant drugs, which are highly addictive and do great harm to society. Studies have shown that the abuse of psychostimulants can cause central nervous system disorders, resulting in changes in the electrical signals in the cerebral cortex (Ersche et al., 2012). In reality, it is difficult to collect acute electroencephalogram (EEG) data of stimulant abuse because drug users often take a mixture of drugs in secret and are closed-off managed after arrest. Meanwhile, the mice experiments can control the variables and compare the EEG changes before and after acute abuse that is more conducive to quantitative analysis (Chang, Kuo, & Chan, 1994; Páleníček et al., 2013). Acute EEG signals are often studied by injecting mice with illegal psychostimulant drugs for five consecutive days to construct an experimental model of acute drug abuse (Brown et al., 2011; Kiyatkin & Smirnov, 2010; Newton et al., 2003). (Chang, Kuo, Tsai, Chen, & Chan, 1995) and (Matějovská, Bernášková, & Šlamberová, 2014) found that the EEG of rats in the abuse group changed greatly in the frequency domain through the experimental paradigm of acute cocaine and methamphetamine abuse. With the rapid development of EEG technology and the evolution of signal processing algorithms, artificial feature sets (Jana, Praneeth, & Agrawal, 2021) extracted from EEG are extended and enriched. Furthermore, multi-domain feature sets can be constructed to analyze the comprehensive response of EEG. Owing to abundant artificial feature sets, plenty of studies have successfully detected mental diseases through machine learning algorithms (Jana et al., 2021). Khajehpour (Khajehpour et al., 2019) calculated the brain functional connection network parameters in the frequency domain to classify the EEG signals of the subjects in the cut-off period of methamphetamine through support vector machine (SVM). In recent years, with the rapid development of deep learning (Bizopoulos, Lambrou, & Koutsouris, 2019; Covert et al., 2019), more and more studies have been conducted to obtain the automatic digital features of EEG through models, which do not need solid theoretical knowledge and complicated calculation like artificial data features to easily obtain more subtle weak features. In addition, it can skillfully learn and integrate multi-domain knowledge automatically through multi-layer input.

In this study, the quantitative EEG characteristics of acute illicit psychostimulants cocaine and methamphetamine abuse were analyzed by multi-domain, including time, frequency, complexity, dynamics, independent domain and a new proposed specific frequency domain. Compare with traditional machine learning algorithm, like SVM, k nearest neighbor (kNN), random forest (RF), deep learning algorithm LSTM-FCN and ResNet50 were input into sequence and matrix feature forms respectively to detect the accuracy of illicit psychotropic substance abuse, in which the efficient time-frequency characteristics were integrated.

## 2 Materials and Methods

### 2.1 Data Collection and Pre-processing

#### 2.1.1 Acquisition Modality

For EEG and electromyography (EMG) recording, four holes were drilled on the mouse skull over the frontal and parietal cortices, and two-channel tethered EEG electrodes were anchored onto the skull while raw EEG obtained by differential signal acquisition method. Simultaneously, two EMG electrodes were inserted bilaterally into the neck muscles, that raw EMG obtained by differential method to determine the mice sleeping time as well as motion time. Electrodes were fixed with dental cement. After the surgical procedures, mice were allowed to recover for at least one week in their home cages before being subjected to the EEG recording and drug abuse experiments. The EEG signals were recorded by standard EEG electrodes and analyzed via LabChart software (ADInstruments). As shown in Figure 1, two tethered EEG electrodes were anchored onto the skull, on account of original single-channel EEG collecting from the differential signal between those two electrodes. Meanwhile, the single-channel EMG was also obtained by the differential signal of two EMG electrodes. EMG signals were used as supplement information to remove the sleeping-state of EEG.

**FIGURE 1.**
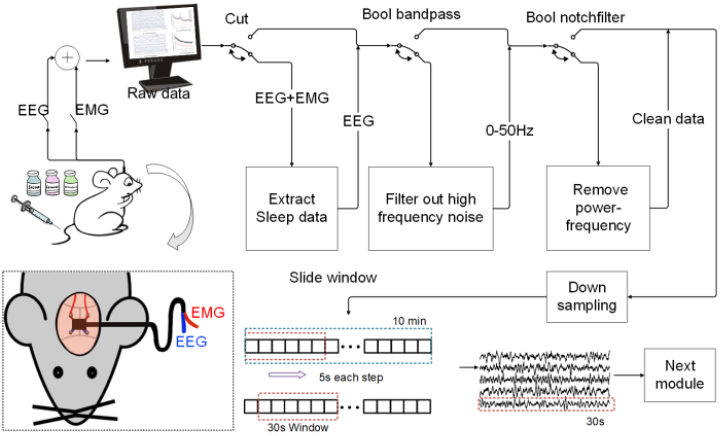
The pipeline of data collection and data pre-processing. Single channel EEG and EMG in resting state were collected, and EEG during sleeping and exercising were removed by referring to EMG. The EEG of 0.1-50 Hz was obtained by band-pass filtering, the power supply frequency interference (50 Hz at P.R. China) was removed by notch filtering, and then sampling was lowered from 1,000 Hz to 200 Hz. The last, data were segmented by sliding window to form a sample and data size are moderate.

#### 2.1.2 Animal subjects and Drugs

Male C57BL/6J mice of 6–8 weeks old were maintained with ad libitum food and water in standard animal rooms with controlled temperature (22 ± 1 ºC) and relative humidity (55% ± 5%) on a 12 h light/dark cycles (lights on from 07:00 to 19:00; 07:00, zeitgeber time 0). The mice were allowed to acclimatize to the recording chambers for at least 3 days before drug treatment and signals recording. All experiments were conformed to the Guidelines for the Care and Use of Laboratory Animals of Zhejiang University, and all protocols were sanctioned by the Institutional Animal Care Committee (No. ZJU20200129). Coca was obtained from Qinghai Pharmaceutical Factory (Qinghai, China). Meth was provided by National Institutes for Food and Drug Control (Beijing, China). Both Coca and Meth were dissolved in 0.9% Sali. During the experiments, Coca (15 mg/kg), Meth (1.0 mg/kg), or Sali (10 ml/kg) were injected intraperitoneally (i.p.) (Dobbs et al., 2016; Kourrich, Rothwell, Klug, & Thomas, 2007). 8 mice were injected with Coca, 8 mice with Meth, and 8 mice with Sali. Relevant drugs were injected every night at 8 o’clock for 5 consecutive days to design the psychostimulants abuse experiment paradigm (Dobbs et al., 2016; Itzhak & Martin, 2000). Awake EEG data were intercepted for about 10 mins after injection and EEG signal was collected at a sampling rate of 1,000 Hz.

#### 2.1.3 Date pre-processing

Refer to Figure 1 for the data collection and pre-processing pipeline. EEG and EMG were collected simultaneously, and EEG during sleeping were removed by referring to EMG signals. Then the waking-state EEG signals in the frequency range of 0.1-50 Hz were extracted through band-pass filtering, which removed the 49.9-50.0 Hz power frequency interference (50 Hz at P.R. China) by a notch filter, then data size was reduced by down-sampling method from 1,000 Hz to 200 Hz. As such, a relatively clean and high-quality EEG data were obtained. Figure 2A-C respectively display power spectrum of mice’ EEG which injected Sali, Coca, and Meth. Since increasing EEG data samples by sliding window segmentation could boost detection performance, we employed a 30s sliding window and 5s sliding step to obtain 13,800 samples for the analysis ultimately. Figure 2D-F respectively display 30 s windows, 5 s overlap EEG’ power spectrum of mice which injected Sali, Coca, and Meth in five days.

**FIGURE 2.**
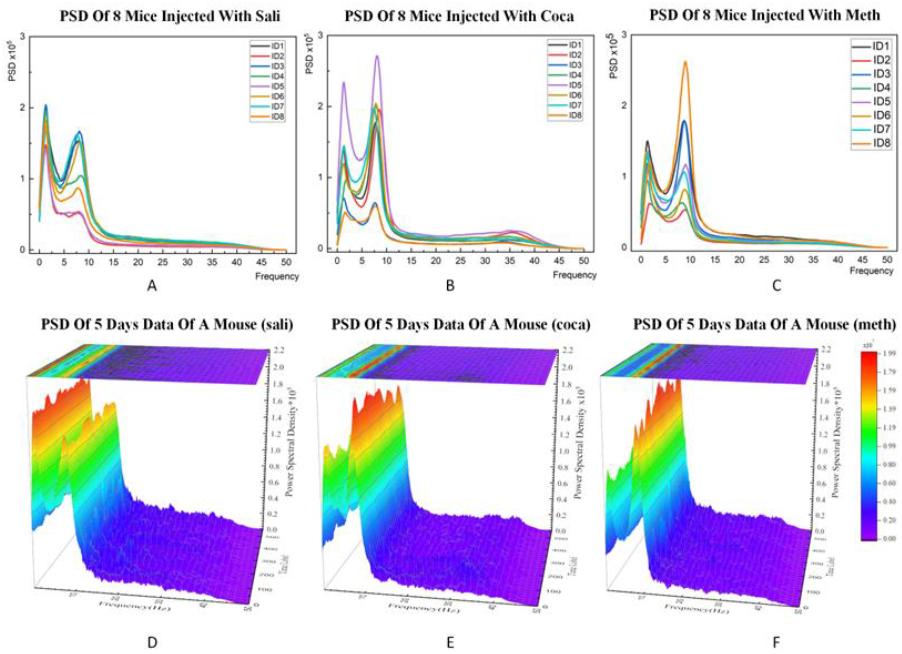
**A-C** respectively display power spectrum of mice’ EEG which injected Sali, Coca, and Meth. **D-F** respectively display 30 s windows, 5 s overlap EEG’ power spectrum of mice which injected Sali, Coca, and Meth in five days.

### 2.2 Feature Extraction

#### 2.2.1 Time Domain

We selected the sum of absolute first difference (FD), variance (Var) and hjorths (Hjorth, 1970) as time domain features. They are commonly used in EEG analysis, on account of FD and Var can show the fluctuations and distribution of signal. Moreover, Hjorth is also diffusely accepted in physiological signal, which defines three time domain features: activity (H_a), mobility (H_m), and complexity (H_c). H_a is the power variance of EEG signal, which measures the degree of deviation of the signal amplitude. H_m is the ratio of the root mean square of the slope of the EEG signal to the root mean square of amplitude, which measures the change of the slope. H_c is the average of the ratio of the slope change to the ideal curve, and the square root is often used to estimate the bandwidth of a signal and measure how many standard slopes there are on an amplitude. The formulas for FD, Var, H_a, H_m and H_c are shown in Supplement formulas 1.

#### 2.2.2 Complexity Domain

Entropy reflects the confusion of EEG in various dimensions, and has a certain correlation with the alertness, reaction ability and neurological disorders of the brain. We selected binned entropy (BinEn) (Skapura & Dong, 2015), approximate entropy (ApEn) (Guo, Rivero, & Pazos, 2010), sample entropy (SpEn) (Jie, Cao, & Li, 2014), permutation entropy (PeEn) (Eckmann, Kamphorst, & Ruelle, 1987), and differential entropy (DE) (Duan, Zhu, & Lu, 2013) for the analysis, as they have high extraction efficiency and discrimination in drug abuse detection. Supplement formulas 2 shows the formulas for BinEn, ApEn, SpEn, PeEn and DE.

#### 2.2.3 Dynamics Domain

Recurrence plot (RP) is a qualitative analysis method of nonlinear dynamic system proposed in 1987 (Zhang & Li, 2014). It is widely used in the analysis of EEG, EMG, and electrocardiogram (ECG) signals (Torse, Khanai, & Desai, 2019). Its essence is to map the high-dimensional motion state trajectory that the naked eye cannot directly judge to the two-dimensional graph, so that the dynamic behavior can be seen intuitively (Li, Ye, Huang, Wang, & Malekian, 2018). Recurrence quantification analysis (RQA) (Webber & Zbilut, 1994) of the nonlinear dynamic system that has been transformed into a recursive graph, then recurrence rate (RR), determinism (DET), laminarity (LAM), averaged length of diagonal lines (L), trapping time (TT) from RP plot are extracted. The formulas for RR, DET, LAM, L and TT are presented in Supplement formulas 3.

#### 2.2.4 Independent Domain

Independent component analysis (ICA) (Koldovský, Málek, Tichavský, Deville, & Hosseini, 2009), in signal processing, is a calculation method used to separate multivariate signals into additive subcomponents. This is done by assuming that the subcomponents are non-gaussian signals and are statistically independent of each other. Extract 30 independent components from EEG origin signal through ICA, and select the top five contributions as the features of the field. Let random EEG data vector *X* = [*X*_1_, *X*_2_, …, *X*_*r*_]^*T*^, where *r* component are mixtures of *r* independent components of random vector *M* = [*M*_1_, *M*_2_, …, *M*_*r*_]^*T*^. Now vector *E* can be expressed as *X* = *AM*, where *A* is a *r* × *r* mixing matrix. The objective of ICA is to find an inverse matrix. Now, independent components *I* can be calculated using *I* ≅ *M* (Jana et al., 2021). At last, we take out the top 5 of the 30 independent component Ica1, Ica2, Ica3, Ica4, Ica5.

#### 2.2.5 Frequency Domain

The five classic rhythms: delta (*δ*), theta (*θ*), alpha (*α*), beta (*β*), and gamma (*γ*), are each related to the various functions of the brain in the frequency domain. The dominant component in EEG changes when the brain performs different functions. Therefore, they are widely used in the identification and classification of EEG signals (Burleigh, Griffiths, Sumich, Wang, & Kuss, 2020; De Aguiar Neto & Rosa, 2019; Fitzgerald & Watson, 2018; Simpraga et al., 2017). So, we extract five classic rhythms in the frequency domain: *δ* (1-4 Hz), *θ* (4-8 Hz), *α* (8-13 Hz), *β* (13-30 Hz) and *γ* (30-50 Hz) by welch’s power spectral density (PSD). Welch method is the best compromise method between averaging and smoothing, and the power spectrum estimation using overlapping windowing is usually better than that using non-overlapping windowing, and the spectrum curve is smoother. Welch estimate one-sided PSD of real-valued EEG, and *X* is divided into the longest possible segments to obtain as close to but not exceed 8 segments with 50% overlap. Each segment is obtained by a hamming window *w*(*n*) = 0.5 + 0.5*cos*(2*π*(*n*/*M*)), 1 < *n* < *M*. *M, L* are windows length and the number of fragments, so *N* = *ML*(*N* < *length*(*X*), since if it can’t divide *n* into an integer number of segments with 50% overlap, *X* is truncated accordingly). Then periodograms are averaged to obtain the PSD estimate. Calculation formula is as follows:

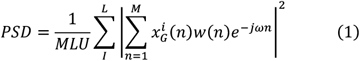

where *U* is the normalizing factor, *ω* is frequency, and [*a, b*] is the band of each rhythm.

#### 2.2.6 Specific Frequency Domain

In the process of original EEG analysis, it finds that comparing with the experimental group and the control group the feature’s value varies greatly in the frequency domain, drug abuse would lead to decrease in the power spectrum, and there was an obvious right shift in the power spectrum in the low frequency band in rapid drug abusing, as shown in Figure 3A, which is consistent with the experimental results obtained in the existing research (Ferger, Kropf, & Kuschinsky, 1994; Ross, Reisfield, Watson, Chronister, & Goldberger, 2012). In addition, we can see that PSD distribution of all mice is concentrated in 0.1-15 Hz, and two peaks appear in delta (1-4 Hz) and theta+alpha (4-13 Hz) frequency bands. We present the mean and standard deviation of five classical rhythms in Figure 3B&C to explore the differences and characteristics in frequency domain between drug abuse addiction and control rats’ EEG. We analyze and extract the characteristics of these two frequency bands. L_S means *PSD*(*delta*) divide *PSD*(0.1 − 49.9*Hz*). L_M means max value in delta band divide *PSD*(*delta*). H_S means *PSD*(*alpha*) divide *PSD* (0.1 − 49.9*Hz*). H_M means max value in alpha band divide *PSD(alpha)*. Diff is equal to the max value in delta minus the max value in alpha. Re_diff is equal to L_M divide H_M.

**FIGURE 3.**
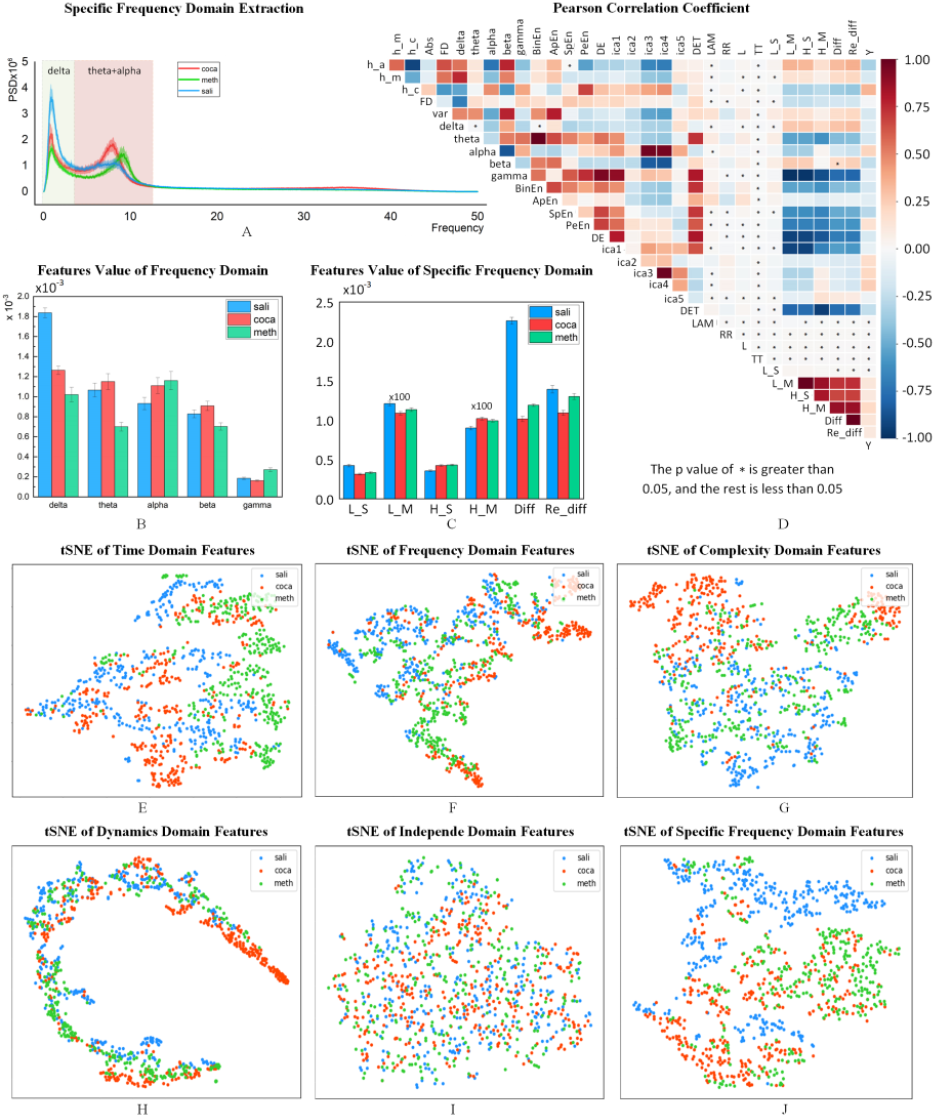
Multi-domain Features Statistical analysis. **A** Average and std of 8 mice’s PSD are shows in shadow region. **B** presented the mean and std of five classical rhythms in frequency domain. **C** presented the mean and std of six features in specific frequency domain. **D** Pearson correlation coefficient between 31 features and category, and grids fill with ‘∗’ means p > 0.05, and the rest is less than 0.05. **E-J** Coca, Meth and Sali subjects after tSNE respectively of time, frequency, complexity, dynamics, independent, and specific frequency domain features.

Where *PSD*(*freq*) is the power spectrum density value between *freq* frequency range. Those six features show a significant difference between the experimental groups and the control group, but not between Coca and Meth. Specific frequency domain shows smaller discrepancy between Coca and Meth in Figure 3C, so as to it performs better in binary detection than in multi-class detection.

### 2.3 Multi-domain Features Statistical Analysis

#### 2.3.1 Quantitative Analysis of Drug Abuse

Six domain features’ value in three groups are demonstrated in Supplement Table 1. Due to the large numeric difference between features caused by the calculation methods of different domains, it is not conducive to using the data of features directly for the detection of drug use by the classifier in machine learning. Therefore, all features are zero-mean normalization first. The contents of ‘(…)’ in the grouping statistical analysis table are negative, and those with ‘e∗’ are 100 times larger than the mean value. Among them, since the small-scale amplitude or the nuance between samples, features in dynamic domain (except the DET) and L_S in specific frequency domain result in smaller mean value than other features. Figure 3 show the statistical analysis of multi-domain features. It is noteworthy that the multiple of DET in dynamics domains is large, resulting in banded distribution of Figure 3H.

**TABLE 1.** Psychostimulants detection result by machine learning

| Task | Method | SVM |  |  |  | RF |  |  |  | kNN |  |  |  |
| --- | --- | --- | --- | --- | --- | --- | --- | --- | --- | --- | --- | --- | --- |
|  |  | Acc | Pre | Sen | F1 | Acc | Pre | Sen | F1 | Acc | Pre | Sen | F1 |
| Binary detection | S_T | 66.52 | 66.69 | 61.47 | 63.97 | 69.87 | 71.94 | 69.87 | 70.89 | 66.59 | 66.61 | 68.08 | 67.34 |
|  | S_F | 73.69 | 76.57 | 69.14 | 72.67 | 73.92 | 75.25 | 73.92 | 74.58 | 70.81 | 72.07 | 70.81 | 71.43 |
|  | S_C | 70.98 | 73.70 | 87.92 | 80.18 | 71.29 | 73.02 | 71.29 | 72.14 | 70.87 | 72.40 | 70.87 | 71.63 |
|  | S_D | 66.49 | 67.37 | 55.24 | 60.7 | 68.88 | 70.19 | 68.88 | 69.53 | 68.93 | 69.22 | 75.00 | 71.99 |
|  | S_I | 63.58 | 63.62 | 62.33 | 62.97 | 64.10 | 64.24 | 64.10 | 64.17 | 63.22 | 63.23 | 64.57 | 63.89 |
|  | S_SF | 74.37 | 74.56 | 78.42 | 76.44 | 74.61 | 75.81 | 74.61 | 75.21 | 71.07 | 72.80 | 82.22 | 77.22 |
|  | M_5V | 75.97 | 75.99 | 77.33 | 76.65 | 74.42 | 74.43 | 75.27 | 74.85 | 73.76 | 73.85 | 70.59 | 72.18 |
|  | M_5M | 76.65 | 76.65 | 75.99 | 76.32 | 78.15 | 79.57 | 78.15 | 78.85 | 76.65 | 76.65 | 75.99 | 76.32 |
|  | M_6V | 78.48 | 78.50 | 79.79 | 79.14 | 79.22 | 80.57 | 88.14 | 84.19 | 78.08 | 78.19 | 90.53 | 83.91 |
|  | M_6M | 78.23 | 78.13 | 83.27 | 80.62 | **81.38** | **81.16** | **88.51** | **84.68** | 79.73 | 79.77 | 81.61 | 80.68 |
| multi-class detection | S_T | 63.95 | 64.10 | 69.13 | 66.52 | 63.55 | 63.03 | 72.85 | 67.59 | 62.89 | 62.93 | 65.55 | 64.21 |
|  | S_F | 65.05 | 65.10 | 68.02 | 66.53 | 69.13 | 70.47 | 69.13 | 69.79 | 62.93 | 63.28 | 71.05 | 66.94 |
|  | S_C | 61.26 | 61.51 | 60.46 | 60.98 | 63.85 | 63.82 | 60.78 | 62.26 | 61.74 | 61.76 | 48.23 | 54.16 |
|  | S_D | 61.25 | 61.25 | 60.85 | 61.05 | 65.40 | 65.82 | 52.23 | 58.24 | 59.77 | 59.61 | 49.48 | 54.07 |
|  | S_I | 51.53 | 51.93 | 74.36 | 61.15 | 57.36 | 57.39 | 60.80 | 59.05 | 59.77 | 59.61 | 49.48 | 54.07 |
|  | S_SF | 66.64 | 66.87 | 72.45 | 69.55 | 70.12 | 74.61 | 48.76 | 58.98 | 69.09 | 69.14 | 71.71 | 70.4 |
|  | M_5V | 69.18 | 70.71 | 79.64 | 74.91 | 71.49 | 71.57 | 74.45 | 72.98 | 70.12 | 74.61 | 48.76 | 58.98 |
|  | M_5M | 64.86 | 66.51 | 63.42 | 64.93 | 66.81 | 66.84 | 66.16 | 66.5 | 65.47 | 65.4 | 59.88 | 62.52 |
|  | M_6V | 74.74 | 74.75 | 75.65 | 75.2 | 73.49 | 73.49 | 74.18 | 73.83 | 74.92 | 75.04 | 71.4 | 73.17 |
|  | M_6M | 75.03 | 75.03 | 75.28 | 75.15 | **76.59** | **76.59** | **77.17** | **76.88** | 76.48 | 76.48 | 76.3 | 76.39 |

#### 2.3.2 Features Correlations

Correlation analysis is conducive to inspect the similarity and the redundancy between features. Therefore, we calculate pearson correlation coefficient between 31 features to demonstrate the relationship inter and intra multi-domain as well as interrelationship and redundancy in feature vectors. We find out feature’s correlations within groups are generally greater than between groups, hence there is valid and complementary information between the domains instead of surplus extraction of numerous features. Dynamics domain have very little correlation with other domains as well as very little similarity among features. Nonetheless, the correlation between frequency, complexity, new frequency domain are strong. Pearson correlation coefficient between 31 features and category is showed in Figure 3D, and grids fill with ‘e∗’ means p > 0.05, and the rest is less than 0.05.

#### 2.3.3 Drug Abuse Difference in Multi-domain

In the above linear correlation analysis, the linear correlation of each domain is various, so t-distributed stochastic neighbor embedding (tSNE) is becomingly adopted to compress the high-dimensional data into a two-dimensional space and verify the nonlinear difference of features in each domain to each category intuitively. The distribution of Coca, Meth and Sali subjects’ feature vectors of each domain after dimensionality reduction visualization was also drawn in Figure 3E&J. These three categories can be distinguished to some extent in time, complexity, frequency, and specific frequency domain. As for dynamics domain (Figure 3H), the large span difference in the two dimensions is caused by the fact that DET is much larger than other parameters. Meanwhile, we find that independent domain is hard to tell them apart (Figure 3I), since sample points of Coca, Meth and Sali are highly mixed.

### 2.4 Multi-domain Detection by Machine Learning

We used SVM, kNN, RF algorithms to detect drug abuse through the features of the six domains mentioned above. Each classifier’s detection result is averaged for the 10 sub-experiments in binary detection task and multi detection task, including 6 single domain classifications, merging 5 domains with the exception of specific frequency domain, merging all 6 domains, using voting mechanism instead of merging for 5 domains and 6 domains tasks. All the features are zero-mean normalized so that the features are in the same dimension. After processing, the mean value of data is 0, and the standard deviation is 1. In machine learning algorithms, datasets are standardized for training to the extent that classifiers have the same emphasis on each feature, which is suitable for occasions with enough samples and can well adapt to noisy EEG data than min-max normalization. To explore effective domain of psychostimulants abuse detection and improve classification effectiveness through single domain detection, multi-domain hybrid detection, and vote-based detection, we take ten-fold cross validation and grid search parameters to get the optimal value, the pipeline of machine learning to detect is show in Figure 4A.

**FIGURE 4.**
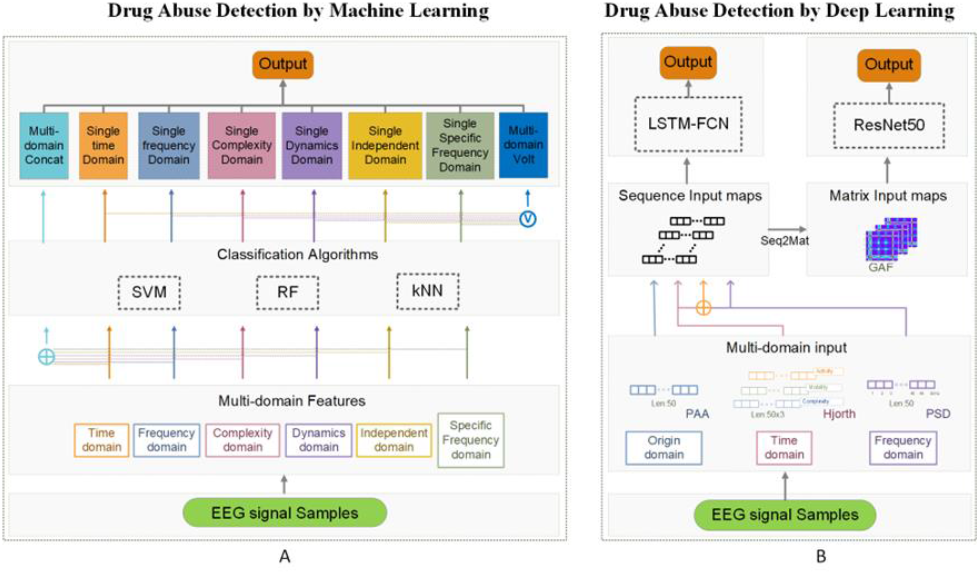
Pipeline of drug abuse detection. **A** Pipeline of drug abuse detection by classic machine learning. From EEG extract out multi-domain features, then we can put each domains into svm, rf and kNN to detect output single time domain detect(S_T), single frequency domain detect(S_F), single complexity domain detect(S_C), single dynamics domain detect(S_D), single independent domain detect(S_I), single Specific frequency domain detect(S_SF), also can concat multi-domain features to one vector to classification algorithm (M_M), or Vote output for single domain output results(M_V). **B** Pipeline of drug abuse detection by deep learning. From EEG extract out multi-domain features(in sequence), then put into Resnet50 after GAF(change sequence to matrix) or put into LSTM-FCN directly.

#### 2.4.1 SVM

The principle of SVM (Wang, 2005) is to find the best separation hyperplane in the data feature space (two-dimensional or multidimensional space), make the maximum interval between positive and negative samples on the training set to correctly divide positive and negative samples. SVM is one of the most widely used supervised learning algorithms, and it can also be used to solve nonlinear problems with kernel method. In this experiment, the grid search method was adopted to select the best hyperparameters of learning rate, gamma, and C. The learning rate parameter list was 0.0001, 0.001, 0.01, 0.1, 0.2 and 0.3, the gamma parameter list was 0.001, 0.01, 0.1 and 1, and C parameter list was 1, 2, 4, 8, 16, 32 and 64.

#### 2.4.2 kNN

kNN represents the unknown sample with the category of its nearest k value, which is one of the simplest classification algorithms (Zainuddin, Lee, Mansor, & Mahmoodin, 2016). In the study of drug abuse recognition and judgment of EEG, whether subjects abuse psychotropic drugs or not can be dichotomized by processing EEG signals and using extracted features as the basis to classify *k* adjacency. In the *k* proximity algorithm, the size of parameter *k* has a decisive influence on the accuracy of the experimental results, so the cross-validation method is selected in this experiment to find the optimal parameter. In addition, the distance measurement of k proximity algorithm has a great influence on the accuracy of experimental results. We use grid search to select the best hyperparameters of weights, neighbors, and p. Weights parameter list is uniform and distance, the values of neighbors are between 0 and 10, and p is between 1 and 5.

#### 2.4.3 RF

RF developed by leoBreiman proved to be an effective ensemble learning method for classification (Vaid, Singh, & Kaur, 2015). In the process of decision tree-based learner, the random variable selection method is introduced on the basis of bagging ensemble. RF uses bootstrap sample technology to establish a set of classification trees, while RF uses multiple trees to train and predict the samples, which is easy to realize and has low computational cost. Besides, due to its diversity (self-sample disturbance and self-attribute disturbance make it “diverse”), in the analysis and recognition task of psychostimulants abuse based on EEG signal, it showed good generalization ability. The grid search method was adopted to select the optimal hyperparameters of decision tree n estimators, max depth, min samples split, and min samples leaf. The n estimators parameter was between 0 and 30, the max depth parameter is between 0 and 5, the min samples split parameter is between 0 and 30, and the min samples leaf parameter is between 0 and 30.

### 2.5 Multi-domain Detection by Deep Learning

Different from classic machine learning to extract plenty of features, deep learning allows us to input larger and more abstract features. Pipeline of classic machine learning and deep learning for drug abusing detection are showed in Figure 4A & B, respectively. EEG is a kind of time series with relatively weak, stable and regular signals. Different from ERP component analysis, the collection of resting EEG needs to last for a certain amount of time to provide enough information, so the sequence length is long, resulting in large model input and corresponding training duration. The sample data in this experiment are all 30-second resting EEG data with a sampling frequency of 200 Hz, namely, time series data with 1 channel and 6,000 length. This study used and compared the three kinds of data preprocessing ways, piecewise aggregate approximation (PAA), Hjorth, and PSD, to preprocess the original EEG signals. Pre-processing can not only artificially extract the internal information that is not obvious in the EEG to help the model training and detection, but also reasonably compress the input of the original 6,000 data points, making the input magnitude more suitable for the model input, reducing the training and testing time, and improving the detection result.

#### 2.5.1 PAA

PAA is a common and effective data dimension reduction method, which can smooth the time series and maintain the trend characteristics of the series for the time series *X* = {*x*_1_, *x*_2_, …, *x*_*n*_} to *m*(*n* > *m*) long sequence = {*q*_1_, *q*_2_, …, *q*_*m*_}.

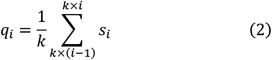

where 1 ≤ *i* ≤ *m, k* = *floor*(*n*/*m*).

#### 2.5.2 Hjorth

Referring to the calculation formula of activity, mobility and complexity in the hjorth parameter in Supplement Formulas 1(Equation 3, 4 and 5), we collected a group of activities at every 120 points in the 6000-point time series, resulting in a set of matrix data of 50 × 50 × 3.

#### 2.5.3 PSD

It can be seen from the above original EEG signal analysis and multi-domain features analysis that in detection of drug abuse, frequency domain information always stands out in distinguishing each category to the greatest extent as well as has a relatively solid theoretical basis. Therefore, we use frequency domain information in deep learning to expand the detection method. Refer to the extraction method of absolute power spectral density parameter of PSD parameter in Equation 1. Extract the power value of each frequency between 0.1-49.9 Hz from the time series of n=6,000, to obtain *F* = {*f*_1_, *f*_2_, …, *f*_50_}. Deep residual learning (ResNet) (He, Zhang, Ren, & Sun, 2016) is a network with deeper depth than general convolutional neural network Net, directly introduces the data output of a previous layer to the input part of the later data layer by skipping multiple layers, which means that the content of the latter layer’s characteristic layer is partly contributed by the linear contribution of a previous layer. Therefore, ResNet50 overcomes the problems of low learning efficiency and low accuracy caused by the deepening of network depth, and has excellent performance and wide application in image classification.

Since pre-processed data are all 50-point sequences (hjorth has three 50-point sequences), which cannot be used as the input data of ResNet50, we need convert one-dimensional inputs into two-dimensional inputs. Comprehensive consideration is taken from various perspectives such as timing relation between data, data fidelity, theoretical support, calculation requirements, etc. In the end, we select the first one in gramian angular fields (GAF) (Oates, 2015), markov transition fields (MTF) (Oates, 2015) and RP (Zhang & Li, 2014). GAF is based on polar coordinates, which retain absolute temporal relationships relative to cartesian coordinates. After converting a scaled time series to polar coordinates, we can easily use the angular perspective to identify temporal correlations over different time intervals by considering the triangular sum between each point. Time series *X* = {*x*_1_, *x*_2_, …, *x*_*n*_}, we change the *X* range so that all values fall in the interval [−1,1]:

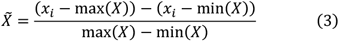

According to this formula, *X* can be expressed in polar coordinates as the time series of rescaling 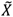 encoding the values as angular cosines and the time stamps as radiuses, as follows:

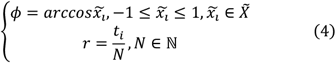

where *t*_*i*_ is the timestamp and ℕ is the constant factor of the span of the normalized polar coordinate system. Over time, the corresponding values are distorted between different points across the circle. GAF is defined as:

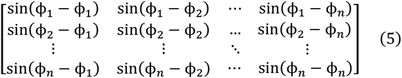

The encoding map of Equation 4 has two important properties. First of all, it’s bijective, because cosine is monotonic at [0, *π*]. Second, relative to the cartesian coordinate and polar coordinate to keep the relationship between the absolute time, from timestamp *i* to the corresponding area of the timestamp *j* depends not only on time interval |*i* − *j*|, but also depends on the absolute value of the *i* and *j*, namely with the position from the top left corner to the lower right corner, time increases.

After converting the scaled time series to a polar coordinate system, we can easily use the angular perspective to identify time correlations over different time intervals by considering the trigonometric sum between each point.

Meanwhile, raw EEG sequence compressed by PAA to 50 - point sequence as origin domain and extracting three hjorth parameters from raw EEG sequence divided into 50 segments to form three 50-point sequences as time domain were selected as the comparative input. So as to sequences from PAA and PSD transform to 1 channel matrixes with size of 50 × 50 in pixels, nevertheless sequences from PAA and PSD transform to 3 channel matrixes with size of 50 × 50 in pixels. In addition to this, we also put forward multi-domain input maps by combine hjorth with PSD matrixes, so that will totally have 4 channel depth input matrixes. Eventually, we put those four kinds of matrixes into ResNet50 model respectively to train a psychostimulants abuse detection model.

As we want to figure out whether the input mode would affect the detection efficiency, we select long short-term memory fully convolutional networks (LSTM-FCN) (Karim, Majumdar, Darabi, & Chen, 2018) as the comparison to ResNet50, which input sequence without GAF transform. Since time domain have three sequences, LSTM-FCN showed outstanding performance in the task of classifying time series signals in the old UCR Time Series Repository (127), which also included EEG data (Karim, Majumdar, & Darabi, 2019). With respect to optimization, the adam algorithm was used to optimize the loss function, and adjust parameters of batch-size and epoch to improve model accuracy.

### 2.6 Performance Metric

Accuracy, Precision, Sensitive, and F1-score are used to evaluate the performance of our drug abuse detector, and were respectively defined as:

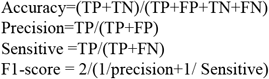

Where TP is the target drug abuser successful detected number; FP is the number of health subjects detected as drug abuser; TN is the healthy subjects successfully detected number; FP is the number drug abuser detected as of health subjects.

Accuracy can demonstrate our detector’s classification ability of each group. Precision shows drug abusing detection ability, when sensitive shows the ability of find out all drug abuser, therefore precision and sensitive are equally important in this task. Owing to consider both of them, F1-score will be paid the closest attention to.

### 2.7 Model Training and Testing

There are two types of detection, first one is binary detection to detect using drug or not, and the second one is multi-class detection which could distinguish between Sali, Coca, and Meth. That means in multi-detection, we used the model to detect not only drug abuse in mice, but also what drugs (Coca or Meth) were injected. We expanded the amount of data through window sliding to meet the training needs. The training duration was 1-30 mins depending on the input size and tasks, and the test duration was less than 0.1s. Training and testing were all implemented on Intel(R) Xeon(R) Silver 4210R CPU@2.40 GHZ GPU RTX3080.

#### 2.7.1 Binary Detection

The classification was based on inter-individual detection, with 12 mice randomly selected as training sets and 4 mice as test sets to detect drug abuse. Among the 16 mice, 8 mice injected Sali were taken as negative examples, and 4 mice injected Coca and 4 mice injected Meth were taken as positive examples, because positive and negative sample equalization can train the model that truly reflects the accuracy rate and sensitivity rate of positive and negative samples.

#### 2.7.2 Multi-class Detection

Used all 24 mice, 8 mice injected Sali, 8 mice injected Coca and 8 mice injected Meth to detect not only drug abuse in mice, but also what drugs (Coca or Meth) were injected in multi-class detection. This ensures not only the balance of data sets but also the maximization of data volume. Same as binary detection of psychostimulants abuse, this task was based on inter-individual detection too.

## 3 Result

### 3.1 Drug abuse detection by Machine Learning

#### 3.1.1 Binary Detection

Table 1 shows the binary classification results obtained by different machine learning algorithm SVM, RF and KNN respectively. The results of single domain detection are all lower than those of multi-domain detection. Among those six domains, frequency domain, complexity domain and specific new frequency domain have better detection result. The poor performance of single domain classifier directly leads to the poor performance of classifier based on voting algorithm. Experimental results show that higher accuracy can be obtained by combining 31 features from these 6 domains. Finally, when max depth is 13, min samples leaf is 3, min samples split is 5, and n estimators is 25, the multi-domain features detection based on RF algorithm achieves the best detection effect with 81.38% accuracy, 81.16% precision, 88.51% Sensitive and 84.68% F1-score, with the results shown in Table S1.

#### 3.1.2 Multi-class Detection

In multi-detection, detections designed by frequency domain and complexity domain are relatively high, which is similar to that of the binary detection. However, the new specific domain is less effective in the multi-class detection because it only distinguishes between drug abusers and non-drug abusers. Similarly, in multi-class detection, the classifier based on voting mechanism performs poorly. Fusion and complementary effect of multi-domain features are better. Any of the six domains can’t accurately distinguish between those three categories, since single domain of EEG characterization of drug abuse is inadequate and one-sided, which leads to poor effect of a single domain detection. Ulteriorly, the voting method based on the detection results of those six domain classifiers is not satisfactory too. As a whole, multi-class detection is harder than the binary detection. Multi-domain features detection based on RF algorithm achieves the best detection result with 76.59% accuracy, 76.59% precision, 77.17% sensitive and 76.88% F1-score, when max depth is 18, min samples leaf is 5, min samples split is 8, and n estimators is 29.

### 3.2 Drug Abuse Detection by Deep Learning

The results are summarized in Table 2. After four methods being used in preprocessing original EEG to produce four domains, detection by deep learning is mainly based on LSTM-FCN and ResNet50 models. Comparing those methods, multi-domain is distinguished in detection, since its input has more comprehensive information. This result is consistent with the emergence of multi-domain features in machine learning. We trained LSTM-FCN and ResNet50 through the grid to optimal parameters.

**TABLE 2.**
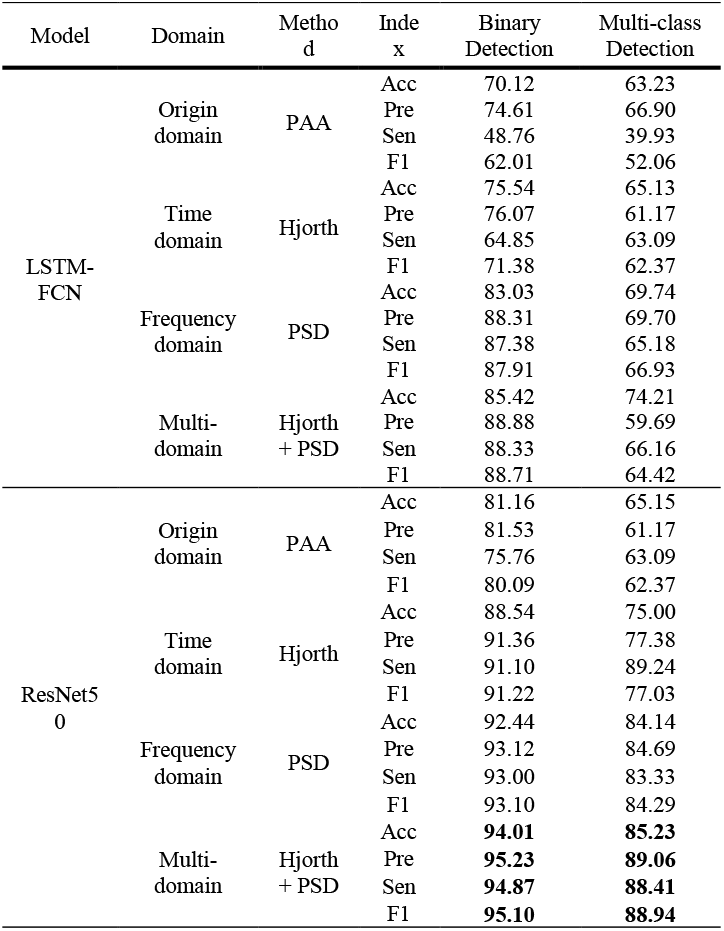
Binary detection and multi-class detection by deep learning

#### 3.2.1 Binary Detection

Original domain performs the worst in either LSTM-FCN or ResNet50, because it extracts valid information from raw EEG data by PAA, which is a technology that gains lightweight input by sacrificing high resolution EEG signals. Meanwhile, consistent with the classification result of machine learning, the classification effect in frequency domain of deep learning is observably better than that in the time domain. The difference is that in the deep learning, the time domain extracts a map with three depths through hjorth, and its input magnitude is three times that of the frequency domain. Learning from multi-domain measure used in machine learning to greatly improves accuracy, combining time domain with frequency domain to produce multi-domain input maps improves detection accuracy slightly. Resnet50 is superior to LSTM-FCN in general. Ultimately, Resnet50 based multi-domain achieves the best binary detection result with 94.01% accuracy, 95.23% precision, 94.87% sensitivity, 95.10% F1-score.

#### 3.2.2 Multi-class Detection

The result of multi-class detection is worse than that of binary detection. Consistent with binary detection, ResNet50 performed better than LSTM-FCN, original domain performs the worst, frequency domain stands out among the three single domains, multi-domain detection performs slightly better than single-domain detection. It can be inferred that in EEG based drug abuse detection scenarios, ResNet50’s ability to learn differences or similarities from multi-channel matrixes is superior to LSTM-FCN’s ability to learn differences between categories directly from time series. At length, multi-domain based ResNet50 achieve the best multi-class detection result with 85.23% accuracy, 89.06% precision, 88.41% sensitivity, 88.94% F1-score.

## 4 Discussion

### 4.1 Disparity in six feature domains

In contrast to numerous previous studies that extracted several features according to specific research purposes, this research extracted comprehensive features from six domains to characterize each category, explored the contributions of each domain as well as the correlationship between features. Meanwhile, multi-domain also improves the accuracy of drug abuse detection. The method allows to detect small differences in waking-state EEG and requires no particular stimulation and task. In the experimental process, we found that the domain based on frequency and complexity had excellent performance. They are also the most frequently selected features of various EEG based analysis. The complexity was based on the disorder degree and entropy of extracted EEG. The disorder degree after drug use was higher than that of the control group. In addition, frequency domain is a widely used method, and the classical rhythm correspondence is relevant to different brain states. Inspired by the frequency domain method, we also added specific frequency domain-based experimental results. Six salient features were added to form the specific frequency domain to improve binary classification, including the specific wave peak and the peak within the frequency band as well as the change of PSD distribution, which provides more comprehensive and suitable frequency domain information for drug detection by EEG. The experimental results show that this domain is helpful only to the binary classification, but is not effective for the multi-class detection.

As for deep learning, frequency domain was directly applied to deep learning by LSTM-FCN based on time series and ResNet50 based on matrix, and the experimental results show that the frequency domain is significantly better than the raw EEG time series origin domain. Meanwhile, results also suggest that properly preprocessed inputs perform better than traditional inputs of raw signals, like frequency and time domain. This also indicates that when deep learning is used to deal with EEG tasks, optimal classification results cannot be obtained only by adjusting model parameters, and effective input preprocessing can obtain unexpected results.

### 4.2 Exceptional detection based on multi-domain method

Because of the abstract, non-intuitive and nuance EEG in this work, the feature set fused with multiple domains has comprehensive information. Furthermore, Large feature sets in the classifier can improve the target of model characterization and prediction, so as to improve the detection accuracy. Experimental results show that multi-domain could significantly improve the detection ability. In machine learning, two multi-domain input methods are commonly used. One method is based on voting mechanism, but its performance is poor due to the limited classification ability of each single domain. The other is based on feature fusion, which integrates all features and significantly improves classification ability. In the binary classification, the accuracy of fusion of 5 domains is lower than that of 6 domains, indicating that more effective features in the machine learning will improve the classification accuracy. Thus, higher accuracy can be achieved by searching for more relevant features. But from the principle of machine learning, it is well known that algorithm will not work well on large data sets, and there will be an upper limit for the data sets. In deep learning, we also conducted multi-domain experiments and obtained the best results. LSTM-FCN and ResNet50 learn more abstract information than machine learning through their deep network models. Furthermore, combine frequency domain information with time domain achieves the best score of drug abuse detection.

### 4.3 Differences between Sequence-based and Matrix-based Model

In deep learning, we mainly relied on the LSTM-FCN and ResNet50 models. LSTM-FCN is based on time series input and learns through the relationship between input sequences. Originally, time and frequency domain can directly input into LSTM-FCN as well as the multi-domain. On the other hand, ResNet50 is based on image input, namely matrix input, and has strong learning ability with a single image. When using ResNet50, we need to convert above data into two-dimensional data through GAF. From input point of view, the input quantity of ResNet50 is the square times that for LSTM-FCN. In this work, 50 × 50 input is used, but the size is within the acceptable range. In terms of testing time, they are very close for the two models. ResNet50 is superior to LSTM-FCN in accuracy. Therefore, the multi-domain drug abuse detection based on ResNet50 model was finally selected, and obtained the best detection results.

### 4.4 Comparison between classic machine learning and deep learning in drug abuse

Machine learning and deep learning were used to identify psychostimulants abuse based on EEG signals. In machine learning, since only slight differences exist among each category, SVM based on maximum interval, kNN based on proximity distance and RF based on decision tree cannot be detected efficiently in 6 domains and 31 features, only 84.68% and 76.88% F1-score respectively in binary and multi-class detection. Therefore, on the basis of previous machine learning results and statistical analysis, we designed deep learning methods for inputting multi-domain input vectors. Ultimately, ResNet50 achieves the highest F1-score: 95.10% in binary detection and 88.94% in multi-class detection. Compared with the detection methods of psychostimulants abuse based on machine learning EEG signals and deep learning EEG signals, machine learning has less training time and higher interpretability of features, but it has lower accuracy. Moreover, feature extraction often requires more professional mathematics, bioelectricity and medical related knowledge. Meanwhile, deep learning has strong learning and generalization abilities, but it requires a large quantity of samples for training and long training time.

### 4.5 Contrast of Binary and multi-class detection

A binary classification task was carried out to study whether the subjects took psychostimulants or not. Meanwhile, the types of psychostimulants being taken were detected as well. Coca and Meth were included in this study, so it was called the multi-class detection task. The classification accuracy of the multi-class detection in the five models of SVM, kNN, RF, LSTM-FCN and ResNet50 are far lower than the binary detection. However, only Coca and Meth were selected in this work, and both of them belong to the psychostimulants, it’s harder to detect out. In the subsequent research, more kinds of drugs are needed for further research.

### 4.6 Future application of EEG based drug abuse

Drug taking will lead to public health and social problems, such as drug driving, epidemic infection and serious crimes. At present, there are few legal detection techniques; the improvement of psychostimulants abuse identification system based on EEG signal could be suitable for drug detection because of its advantages of non-originality, wide availability, relatively low price, excellent record resolution and relatively simple use. However, achieving these goals will require accurate acquisition hardware, portable EEG caps, more durable and conductive electrodes, and better anti-interference circuitry. Meanwhile, expand dataset to reduce individual differences, and then an EEG acquisition and analysis system suitable for spot examination was constructed. Ultimately, a drug driving detection system based on EEG signal can be developed with some improvement, which can assist roadside drug driving detection which is low cost, fast and wide.

## 5 Conclusion

This paper developed an effective drug abuse recognition method and an optimum system. Firstly, 31 features were extracted from the six domains, including time domain, frequency domain, complexity domain, dynamics domain, independent domain, and specific frequency domain. Through the input methods of single domain, multi-domain and volt-based multi-domain, binary detection and multi-detection tasks were carried out by machine learning algorithms SVM, KNN and RF. RF algorithm surpasses others with F1 score of 84.68% and 76.88% respectively in binary detection and multiple detection. As for deep learning, this study explored acute drug abuse detection by LSTM-FCN and ResNet50 for the sake of optimizing the detection system, respectively basing original signals, time domain signals, frequency domain signals, and multi-domain data as input of LSTM-FCN and ResNet50. Finally, the multi-domain sequence data transformed into matrix through GAF and entered ResNet50 to get the best detection result, with F1 score of 95.10% and 88.94% respectively in binary detection and multiple detection. The proposed system and algorithms show promising application in fast field detection for drug abuse in recent future.

## Acknowledgements

This work was supported by grants from the National Key R&D Program of China (2018YFA0701400), Zhejiang Province Key R & D programs (No. 2021C03003, No. 2021C03062 No. 2021C03108).

