## supplemental Table 1 for "Multi-Domain Characterization and Rapid Detection Technology for Cocaine and Methamphetamine Acute Abuse based on EEG": supplement.docx

**S1.** The formulas for FD, Var, H_a, H_m and H_c.

The original EEG signals are defined as $X = \{x(i), i=1, 2, \ldots, n\}$.

$$\begin{aligned} FD = \sum_{i=1}^{n-1} \left| x_{i+1}-x_{i} \right|\#\left( 1 \right) \end{aligned}$$

$$\begin{aligned} Var=\frac{\sum_{i=1}^{n} \left( x_{i}-\mu\right)^{2}}{n-1}\#\left( 2 \right) \end{aligned}$$

$$\begin{aligned} H_{a}=\sigma_{X}^{2}\#\left( 3 \right) \end{aligned}$$

$$\begin{aligned} H_{m}={{\sigma_{X^{'}}}/\sigma}_{X}\#\left( 4 \right) \end{aligned}$$

$$\begin{aligned} H_{c}={\frac{\sigma_{X^{''}}}{\sigma_{X^{'}}}}/{\frac{\sigma_{X^{'}}}{\sigma_{X}}}\#\left( 5 \right) \end{aligned}$$

where $\mu$ is the mean of $X$. $X'$and $X''$ are first difference and second difference respectively. $\sigma_{X}$, $\sigma_{X^{'}}$, $\sigma_{X^{''}}$are the standard deviations of $X$, $X'$, $X''$.

**S2.** The formulas for BinEn, ApEn, SpEn, PeEn and DE.

BinEn was obtained by equidistant box partitioning. By dividing the value of $X$ into boxes, the interval $[min(X), max (X)]$ can be equally divided into *K* bins. Then the value of $X$ will be distributed among the $K$ bins, hence BinEn can shows that the sequence amplitude is evenly distributed or concentrated in some intervals. The entropy of the probability distribution can be calculated as follows:

$$\begin{aligned} BinEn=-\sum_{k=0}^{Bins} p_{k}\log\left( p_{k} \right)\#\left( 6 \right) \end{aligned}$$

where $Bins$ are $min(bins,len(X))$, $p_{k}$ is the percentage of samples in bin $k$, and $bins=10$.

ApEn is a measure of complex types of unstable time sequence, with the view of detecting the probability of new subsequence generation in time series. EEG signal $X$ is reconstructed to m-dimensional vector $\{Y = y(i),i=1, 2, \ldots, M, M = n-m+1\}$, and $y(i) = \{x(i), x(i+1), ... , x(i+m-1)\}$, where $m$ is the embedding dimension and the length of the window (sequence participating in the comparison). The maximum distance between each component is defined as the maximum contribution component distance $D\{y(i),y(j)=max\{\left| y\left( i+k \right),y\left( j+k \right) \right|\}, i,j\in\left[ 1, n-m+1 \right], k\in[0,m-1]$. Given threshold $r$, define the ratio of the number of $D\{y(i),y(j)\}<r$ in $y$ as $C_{i}^{m}(i)$ to compute the probability magnitude measure representing the regularity of the sequence $y(i)$.

$$\begin{aligned} \phi^{m}\left( r \right)=\frac{1}{n-m+1}\sum_{i=1}^{n-m+1} lnC_{i}^{m}\left( i \right)\#\left( 7 \right) \end{aligned}$$

Then calculate $C_{i}^{m+1}(i)$ and $\phi^{m+1}\left( r \right)$ of $m+1$ embedding dimension.

$$\begin{aligned} ApEn=\phi^{m}\left( r \right)-\phi^{m+1}\left( r \right)\#\left( 8 \right) \end{aligned}$$

SpEn is only a small transformation of some steps in ApEn calculation. SpEn have certain independence, consistency, such as optimal point. Theoretically SpEn has higher accuracy and efficiency than ApEn, due to solve ApEn's problem of relying on their own data length flawed. SpEn is a method used to measure the complexity of time series, like ApEn, which has been applied in assessing the complexity of physiological time series and diagnosing the state of cases. Whereas they have same steps of reconstruct to m-dimensional embedding $Y$, the distance $D\{y(i),y(j)\}$ of SpEn is defined as the absolute value of the maximum difference between $y(i)$ and $y(j)$, $D\{y(i),y(j)=max\{\left| y\left( i+k \right)-y\left( j+k \right) \right|\}$, $i, j\in$ $\left[ 1,n-m+1 \right], k\in[0,m-1]$. Given threshold $r$ and embedding dimension $m$, define the ratio of the number of $D\{y(i),y(j)\}<r$ in $y$ as $B^{m}(r)$ to compute the probability magnitude measure representing the regularity of the sequence $y(i)$.

$$\begin{aligned} A^{m}\left( r \right)=\frac{1}{n-m}\sum_{i=1}^{n-m} B_{i}^{m}\left( r \right)\#\left( 9 \right) \end{aligned}$$

Then calculate $A^{m+1}\left( r \right)$ of $m+1$ embedding dimension.

$$\begin{aligned} SpEn=\ln A^{m}\left( r \right) -\ln A^{m+1}\left( r \right)\#\left( 10 \right) \end{aligned}$$

PeEn algorithm, a method for detecting dynamic mutation and randomness of time series, quantitatively evaluate random noise contained in signal sequences and a dynamic mutation detection, which not only can conveniently and accurately locate the moment of system mutation but also can amplify the small signal. EEG signal *X* is reconstructed to the m-dimensional vector $\{y(i),i=1,2,\ldots,m, M=n-(m-1)t\}$,and $y(i)=\{x(i),x(i+t),...,x(i+(m-1)t)\}$, Where $m$ is the embedding dimension, $t$ is the delay time. Each reconstruction portion is rearranged in ascending order to obtain the column index of each element position in the vector to form a series of symbols, and there are altogether $m!$ different symbolic sequence mappings in m-dimensional phase space. Count the number of occurrences of each sequence divided by the total number of occurrences of $m!$ different symbol sequences as the probability of the reconstruction weight. The probability distribution of all symbols is represented by $\left\{ P_{1},P_{2},...,P_{K} \right\}, K\leq m!$.

$$\begin{aligned} PeEn=\frac{-\sum_{j=1}^{k} P_{j}\ln\left( P_{j} \right)}{\ln\left( m! \right)}\#\left( 11 \right) \end{aligned}$$

DE is the generalized form of shannon information entropy on continuous variables and is widely used in EEG classification tasks. An EEG that approximates a gaussian distribution $N(\mu,\sigma_{i}^{2})$ for a given length, the calculation formula is as follows:

$$\begin{aligned} DE=-\int_{a}^{b} p\left( x \right)\log\left( p\left( x \right) \right)dx\#\left( 12 \right) \end{aligned}$$

where $[a,b]$ is the value range, $p(x)$ is continuous probability density function of characteristic information.

**S3.** The formulas for RR, DET, LAM, L and TT.

RR represents the ratio of black dots in a recursive graph to points in the entire recursive graph. It tells us the proportion of point vectors that are close to each other in the *m*-dimensional phase space. For the time series of $n$ points, RR can be calculated by:

$$\begin{aligned} RR=\frac{1}{N^{2}}\sum_{i=1}^{N} \sum_{j=1}^{N} R_{i,j}\left( \epsilon\right)\#\left( 13 \right) \end{aligned}$$

DET refers to the percentage of recursion points forming diagonals in a small part of the recursion graph (the percentage of black points on the line segment forming parallel diagonals), which reflects the duration of trajectory determination approximation in the phase space of EEG. The longer the approximation time is, the larger the DET indicates the greater the predictability of EEG.

$$\begin{aligned} DET=\frac{\sum_{l=l_{min}}^{N} lp\left( l \right)}{\sum_{i,j}^{N} R_{i,j}}\#\left( 14 \right) \end{aligned}$$

where, $l_{\min}$ is the minimum diagonal length, $l_{\min}=2$, $p(l)$ is the proportion of the diagonal with length $l$.

LAM, the percentage of recursion points that make up vertical/horizontal lines.

$$\begin{aligned} LAM=\frac{\sum_{v=v_{min}}^{N} vp\left( v \right)}{\sum_{v=1}^{N} Rv}\#\left( 15 \right) \end{aligned}$$

where $p(v)$ is the probability distribution density of vertical line length $v$. $v_{\min}$ is minimum analysis length,$v_{\min}=2$.

Averaged length of L can be calculated by:

$$\begin{aligned} L=\frac{\sum_{l=l_{min}}^{N} Nlp\left( l \right)}{\sum_{l=l_{min}}^{N} Np\left( l \right)}\#\left( 16 \right) \end{aligned}$$

TT includes recursive the number and length of vertical structure in RP plot.

**Table S1** Quantitative analysis of multi-domain features

|  | Sali  (n=8,c=4600) | Coca  (n=8,c=4600) | Meth  (n=8,c=4600) |
| --- | --- | --- | --- |
| H_a | 0.594$\pm$1.093 | (0.385) $\pm$0.809 | (0.209) $\pm$0.779 |
| H_M | 0.336$\pm$1.130 | (0.280) $\pm$0.865 | (0.056) $\pm$0.884 |
| H_c | (0.534) $\pm$1.015 | 0.228$\pm$0.875 | 0.307$\pm$0.879 |
| FD | (0.230) $\pm$1.090 | 0.502$\pm$0.864 | (0.273) $\pm$0.828 |
| Var | 0.301$\pm$1.285 | (0.185) $\pm$1.008 | (0.117) $\pm$0.443 |
| Delta | 0.401$\pm$1.109 | (0.520) $\pm$0.746 | 0.119$\pm$0.877 |
| Theta | 0.162$\pm$1.158 | 0.106$\pm$1.119 | (0.268) $\pm$0.546 |
| Alpha | (0.601) $\pm$1.038 | 0.648$\pm$0.861 | (0.048) $\pm$0.631 |
| Beta | 0.484$\pm$1.300 | (0.361) $\pm$0.819 | (0.123) $\pm$0.511 |
| Gamma | (0.111) $\pm$0.825 | 0.457$\pm$1.123 | (0.347) $\pm$0.848 |
| BinEn | 0.162$\pm$1.158 | 0.106$\pm$1.119 | (0.268) $\pm$0.546 |
| ApEn | 0.163$\pm$1.303 | (0.039) $\pm$1.106 | (0.124) $\pm$0.195 |
| SpEn | 0.176$\pm$0.976 | 0.335$\pm$1.128 | (0.511) $\pm$0.608 |
| PeEn | (0.225) $\pm$0.779 | 0.071$\pm$1.002 | 0.154$\pm$1.145 |
| DE | 0.048$\pm$0.909 | 0.299$\pm$1.107 | (0.347) $\pm$0.858 |
| Ica1 | (0.193) $\pm$0.730 | 0.594$\pm$1.292 | (0.401) $\pm$0.496 |
| Ica2 | (0.323) $\pm$1.208 | 0.194$\pm$0.791 | 0.129$\pm$0.869 |
| Ica3 | (0.587) $\pm$1.074 | 0.664$\pm$0.794 | (0.077) $\pm$0.652 |
| Ica4 | (0.606) $\pm$1.093 | 0.607$\pm$0.803 | (0.001) $\pm$0.653 |
| Ica5 | (0.110) $\pm$1.201 | 0.351$\pm$0.733 | (0.241) $\pm$0.909 |
| DET | 0.250$\pm$0.845 | 0.105$\pm$1.072 | (0.356) $\pm$0.968 |
| LAM* | (0.008) $\pm$1.000 | 0.116$\pm$1.000 | (0.115) $\pm$1.000 |
| RR* | 3.123$\pm$1.000 | (4.037) $\pm$1.000 | (3.900) $\pm$1.000 |
| L* | 1.267$\pm$1.000 | (6.480) $\pm$1.000 | 2.014$\pm$1.000 |
| TT* | 3.827$\pm$1.000 | 1.028$\pm$1.000 | (1.635) $\pm$1.000 |
| L_S* | (1.750) $\pm$1.000 | (1.771) $\pm$1.000 | (2.827) $\pm$1.000 |
| L_M | 0.115$\pm$0.736 | 0.736$\pm$(0.519) | 0.403$\pm$0.742 |
| H_S | 0.132$\pm$0.662 | 0.662$\pm$(0.527) | 0.395$\pm$0.689 |
| H_M | (0.159) $\pm$0.776 | 0.776$\pm$(0.178) | 0.338$\pm$1.062 |
| Diff | (0.053) $\pm$0.675 | 0.675$\pm$(0.205) | 0.258$\pm$1.386 |
| Re_diff | (0.014) $\pm$0.725 | 0.725$\pm$(0.239) | 0.253$\pm$1.366 |

All features are zero-mean normalized. The contents of '(…)' in the grouping statistical analysis table are negative, and features with ‘*’ are 100 times larger than their real mean value.

$$\begin{aligned} TT=\frac{\sum_{v=v_{min}}^{N} vp\left( v \right)}{\sum_{v=v_{min}}^{N} p\left( v \right)}\#\left( 17 \right) \end{aligned}$$
